# Uncovering uncharacterized binding of transcription factors from ATAC-seq footprinting data

**DOI:** 10.1101/2023.10.26.563982

**Authors:** Hendrik Schultheis, Mette Bentsen, Vanessa Heger, Mario Looso

## Abstract

Transcription factors (TFs) are crucial epigenetic regulators, which enable cells to dynamically adjust gene expression in response to environmental signals. Computational procedures like digital genomic footprinting on chromatin accessibility assays such as ATACseq can be used to identify bound TFs in a genome-wide scale. This method utilizes short regions of low accessibility signals due to steric hindrance of DNA bound proteins, called footprints (FPs), which are combined with motif databases for TF identification. However, while over 1600 TFs have been described in the human genome, only ∼700 of these have a known binding motif. Thus, a substantial number of FPs without overlap to a known DNA motif are normally discarded from FP analysis. In addition, the FP method is restricted to organisms with a substantial number of known TF motifs. Here we present DENIS (**DE N**ovo mot**I**f di**S**covery), a framework to generate and systematically investigate the potential of de novo TF motif discovery from FPs. DENIS includes functionality i) to isolate FPs without binding motifs, ii) to perform de novo motif generation and iii) to characterize novel motifs. Here, we show that the framework rediscovers artificially removed TF motifs, quantifies de novo motif usage during an early embryonic development example dataset, and is able to analyze and uncover TF activity in organisms lacking canonical motifs. The latter task is exemplified by an investigation of a scATAC-seq dataset in zebrafish which covers different cell types during hematopoiesis.

## Introduction

The expression of genes has to be tightly regulated to ensure a swift and efficient reaction to changing environmental conditions. One important part of the regulatory machinery in living cells are transcription factors (TF). TFs are DNA binding proteins that control expression by binding to regulatory regions, such as promoters or enhancers, which regulate target genes^1^. TFs utilize specific protein domains to target selected DNA sequences, thus enabling regulation of very few specific genes as well as large gene sets. These binding domains often translate to a preferred DNA sequence motif that can be used to describe the binding affinity of a particular TF. In this context, computational algorithms intended to generate such motifs (e.g. MEME and GLAM2^2,3^) normally utilize Chromatin Immuno-Precipitation sequencing (ChIP-seq^4^ or CUT&RUN^5^ data to identify one TF binding motif at a time. These experimentally derived motifs are collected and organized in specialized TF databases such as JASPAR^6^ or HOCOMOCO^7^. However, these investigations are hindered by the need for TF-specific antibodies for CUT&RUN or ChIP-seq, which are not available for all TFs, or in all organisms of interest. Thus, while more than 1600 genes in the human genome are characterized to be TFs, less than 50% of these have a validated binding motif (JASPAR CORE vertebrates: 727 human motifs). Of note, human and mouse are some of the most studied models in terms of TF binding, but for most other organisms, the rate of TFs with known motifs is considerably lower (JASPAR CORE vertebrates: 14% non-human).

In contrast to antibody affinity based methods, Assay for Transposase Accessible Chromatin using sequencing (ATAC-seq) allows for the unbiased genome-wide assessment of chromatin accessibility. This is realized by a hyperactive Tn5 transposase, which preferentially cuts and inserts sequencing adapters into open chromatin, resulting in fragments of different sizes. After amplification, sequencing, and mapping of the ATAC-seq fragments, open chromatin regions are identified as peaks. Notably, within these peaks, the distribution of cutsites reveals so-called footprints (FPs), which are defined as small regions of reduced read coverage, resulting from DNA-protein binding that protects DNA from Tn5 cutting. This FP signal allows for the genome-wide investigation of TF binding for all known TFs from one ATAC-seq run, as FPs can typically be assigned to known motifs from TF motif databases^8,9^ (Figure 1a). But in some cases FPs can be observed for which there is no known TF motif. This can be hypothesized to be caused by proteins without a known binding motif or with multiple motifs, of which some are unexplored. We define this concept as the uncharacterized binding motif (UBM). Of note, the amount of UBM’s is expected to differ depending on the organism, tissue and cell type under investigation. Thus, a computational tool for UBM characterization needs to identify and visualize de novo motifs and their corresponding gene targets in a cell type and organism-specific manner.

**Figure 1:**
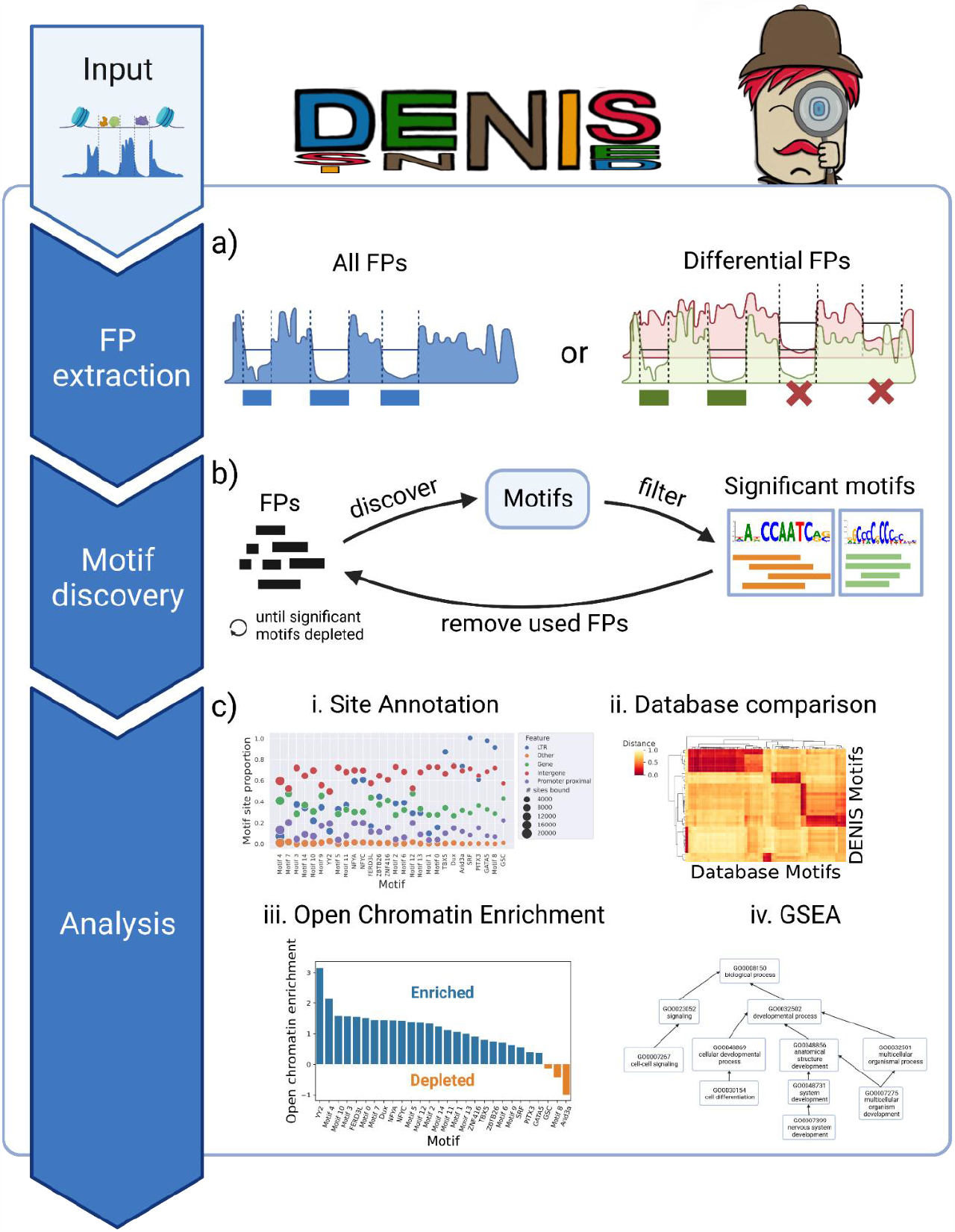
DE Novo motIf diScovery (DENIS) framework. **a)** The framework receives continuous binding scores as minimal input. In the first step, the binding scores are scanned and footprints (FPs) are extracted genome-wide. If binding scores for two conditions are provided as input, differential binding will be assessed. **b)** Sequences at detected FPs are extracted and used to perform data driven de novo motif prediction. In an iterative process, most prominent motifs are extracted and corresponding sequences are removed from the motif discovery pool. FPs not (yet) used are kept for following discovery runs. **c)** Resulting motifs are subjected to downstream analysis. DENIS provides i) an individual motif site annotation module (e.g. annotation to genes), ii) a motif database comparison module, iii) a module to compute global binding site enrichment in open chromatin, and iv) based on the annotation module, a motif based gene set enrichment analysis (GSEA) module. All subfigures were created with BioRender.com

In order to systematically investigate the potential of a genome-wide unbiased FP analysis, we implemented a computational framework called DENIS (**DE N**ovo mot**I**f di**S**covery) that i) isolates UBM events from ATAC-seq data (Figure 1a), ii) performs de novo motif generation (Figure 1b), iii) calculates information content, motif novelty and quality parameters, and iv) characterizes de novo motifs through open chromatin enrichment analysis, differential analysis, gene annotation and gene set enrichment analysis (Figure 1c). The framework is designed to robustly explore DNA binding events on a global scale, to compare ATAC-seq datasets from one or multiple conditions, and is suitable to be applied to any organism. Of note, the latter feature provides potential to use the framework for organisms with very few known TF motifs in order to predict organism-specific TF motifs, and subsequently assign them to conserved TF classes by motif similarity search.

As a proof of principle, we show that DENIS rediscovers the TF Dux in a simulated leave-one-out approach on a bulk ATAC-seq dataset from mouse embryonic stem cells. In addition, the application to a Dux overexpression vs. Dux wildtype (WT) condition revealed a number of differentially regulated de novo motifs indicating several UBMs within early embryonic stages. Finally, the framework is shown to predict both known and de novo TF motifs in a cell type specific manner by the application to a high resolution zebrafish derived single-cell ATAC-seq (scATAC-seq) dataset on hematopoiesis.

## Results

### Workflow and strategy

We implemented the DENIS tool as a modular computational python framework, which smoothly integrates with our previously published software TOBIAS^8^. In order to collect the positions with evidence of protein binding, DENIS firstly detects FPs based on the continuous FP scores from TOBIAS (Figure 1a). This is done by applying a custom peak-calling algorithm to detect stretches of locally increased FP scores. Next, in order to isolate locations of potentially unknown TF binding, DENIS excludes FPs which have a sufficiently good hit for at least one of the motifs included in a reference motif database. In order to consider TF complexes binding in close proximity, DENIS allows for the removal of the motif location within a FP, rather than the complete FP. The remaining FPs are evaluated for their size and location, and FPs below the minimum size threshold are removed. And the genomic sequences within the remaining FP locations are further used for de novo motif generation (Figure 1b). Once the de novo motif generation is finished, UBMs are validated through analysis based on comparison to motif databases and investigation of potential binding locations (Figure 1c). The DENIS framework source code, its documentation and exemplary data is freely available from GitHub.

In order to implement an optimal prediction of motifs (Figure 1b), we investigated three widely accepted tools for de novo motif prediction, namely GLAM2^3^, MEME^2^ and DREME^10^. Through performance tests considering speed, RAM usage, accuracy and precision, we found that MEME was the most suitable motif generation tool to integrate into our application (Supp. Figure 1a-c). However, since MEME was created for the identification of motifs within one ChIP dataset, its application for a mixture of FP derived sequences representing multiple TFs with vastly different input quantities has not been validated. In order to test MEME for this ability, we collected the source ChIP-seq of 20 random motifs originating from JASPAR and constructed a dataset ranging from 395 to 322.803 representative sequences per motif with 659.078 sequences in total. Testing on this set of *in silico* binding sites, we found that MEME was prone to find only the most prominent motifs from a mixture of sequences (Figure 2a).

**Figure 2:**
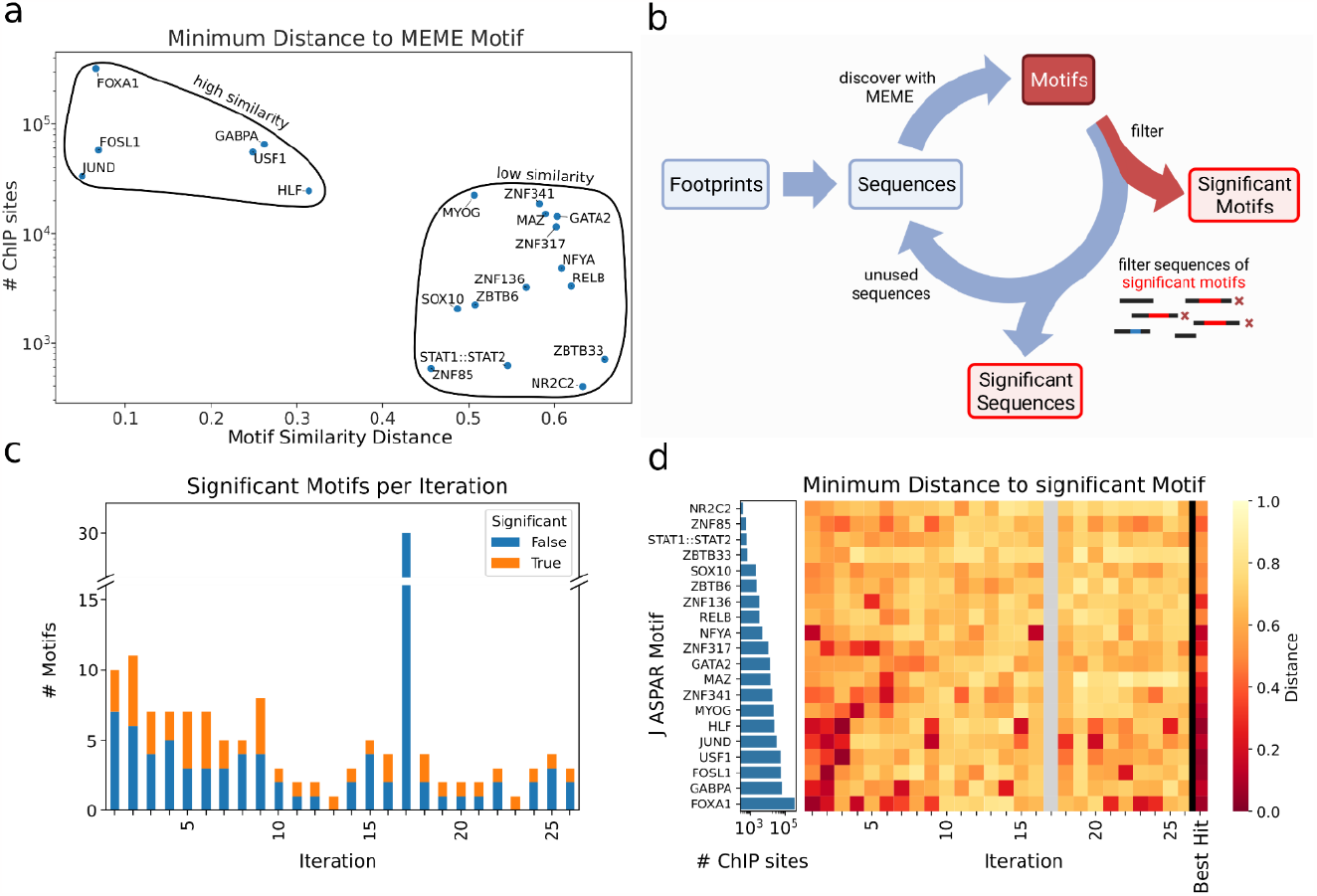
Iterative motif discovery from complex sequence mixtures. **a)** Simulated de novo motif discovery by MEME, each dot indicates the closest MEME created motif compared to the respective original motif; As indicated by black circles, there is a correlation of binding site count (y axis) and de novo motif similarity distance (x axis) to original motif. **b)** Scheme of the iterative motif prediction; Identified FP sequences are extracted to build the motif discovery pool and used to create de novo motifs (red box). Created motifs are filtered for significance and all corresponding sequences used to generate significant motifs are removed from the sequence discovery pool. This process is run iteratively until no further significant motifs are predicted. Figure was created with BioRender.com. **c)** Number (y axis) of generated motifs per iteration (x axis), split by significance (orange and blue) i.e. motifs accepted by internal dynamic e-value filter. **d)** Heatmap showing the distance of de novo motifs detected per iteration (columns) to the most similar database motif (row) for significant motifs from a simulated de novo run. To the left, the number of sites from the corresponding ChIP-seq run per factor is shown as a bar plot. Grey color indicates no significant motifs were available within one iteration.

In order to overcome this resolution issue, we designed DENIS to apply an iterative approach to i) collect the most prominent motifs, ii) filter out the sequences that were used to form these *de novo* motifs, and iii) repeat this process until there are no further significant motifs available in the data (Figure 2b-c). Further filtering steps on the level of motif significance improved the specificity of the framework by reducing the number of motifs while retaining sensitivity (Figure 2d). Finally, DENIS merges very similar motifs found in multiple iterations and continues with the consensus motif in these cases. Using the described test data above, DENIS generated a total of 141 motifs over 26 iterations, which were finally merged to 30 unique motifs. Using this approach, DENIS was able to correctly identify 60% (35% increase to a single run) of the test input motifs. In conclusion, DENIS is capable of finding de novo motifs from a complex *in silico* generated mixture of sequences such as found within uncharacterized ATAC-seq FPs.

To the best of our knowledge, just four other tools exist intended to generate *de novo* motifs from ATAC-seq data, namely BindVAE^11^ and MMGraph^12^ (both using machine learning in combination with k-mers), CEMIG^13^ (which utilizes De Bruijin graphs created on k-mers), and the RSAT peak-motifs pipeline^14^ (a pipeline intended for ChIP-seq, which is also applicable to ATAC-seq data). However, BindVAE, CEMIG and RSAT solely operate on the sequences of complete ATAC-seq peaks, therefore these tools lack precision compared to a FP based tool. While MMGraph utilizes FPs for its de novo motif generation, similarly to DENIS, it does not provide any downstream analysis. As such, these tools can be classified as de novo motif generation tools, similar to MEME, and therefore might be incorporated into DENIS as alternative motif callers in the future.

### Generation of de novo motifs from footprinting tracks

Next, we aimed to utilize ATAC-seq data combined with genomic footprinting for identifying de novo motifs globally within a biological context. For this, we utilized bulk ATAC-seq data of mouse embryonic stem cells (mESC)^15^. Importantly, the data includes two conditions, one with an induced expression of the TF Dux (Dux positive) and a WT condition, where Dux is not expressed (Dux negative). Dux is known to be a major driver for zygotic genome activation during early embryonic development, and its overexpression is sufficient to direct mESCs into a 2 cell-like stage^15,16^. As Dux is not expressed in natural embryonic stem cells, it is not expected to leave any FP in this condition. The motif for Dux should therefore not be discoverable in these cells.

Thus, we asked whether DENIS would be able to specifically identify Dux in the Dux positive cells. As a proof of principle, we excluded Dux and Dux-like motifs (Supp. Figure 2a, b) from the known motif reference database, which enables the Dux motif to be found as part of the UBM’s (Figure 3a), and ran DENIS for both conditions (Dux positive and negative) separately. DENIS found a total of 48 motifs for the Dux positive and three motifs for the Dux negative condition. To verify whether DENIS identified a motif which can be attributed to the Dux factor, we used a previously published ChIP-seq for Dux^15^. The genome was scanned for binding sites, followed by binding site enrichment analysis of ChIP-peaks against the whole genome. We found the highest enrichment for motif 20, which is a motif with a binding profile similar to Dux (Figure 3b). Interestingly, motif 1 of the Dux negative condition was also found among the top enriched motifs. This motif was found to be highly similar to Prdm4, a TF known to regulate pluripotency and differentiation in embryonic stem cells, which explains why it creates FPs in both conditions^17^. Hence, DENIS meets our expectations of recalling biologically relevant motifs from ATAC-seq FPs using a leave-one-out approach as highlighted by rediscovering Dux in the Dux positive condition only.

**Figure 3:**
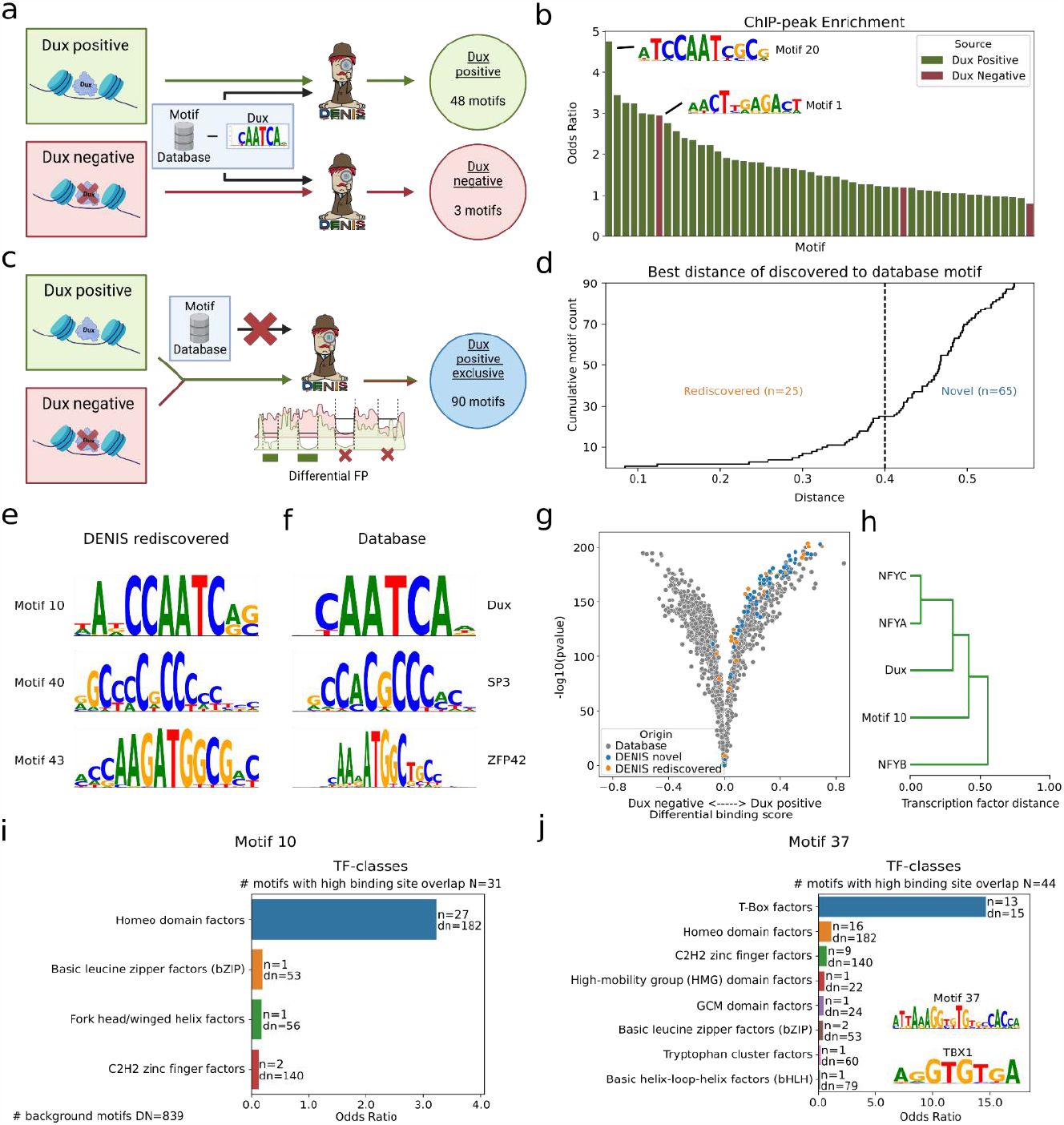
De novo rediscovery of motifs and differential analysis. **a)** Scheme on rediscovering the Dux motif by removal from the background motif database. Separate DENIS runs on a Dux positive and on a Dux negative condition. **b)** Binding site enrichment (y axis) of motifs calculated by DENIS (x axis) as described in (a) within Dux ChIP-seq peaks. The top motifs enriched for Dux positive (green) and Dux negative (red) conditions are shown. **c)** Scheme on how to discover motifs unique to the Dux positive condition without the use of any motif database. DENIS uses the differential FP module to predict motifs exclusively present in the Dux positive condition. **d)** Differential motifs as generated in c), sorted by distance (x axis) to their closest match in the motif database. Motifs below a distance of 0.4 are considered as rediscovered database motifs, motifs above are considered novel. **e-f)** Examples of rediscovered motifs and their corresponding JASPAR database matches. **g)** Combined differential motif activity plot of JASPAR and DENIS de novo motifs. For each motif the fold change of activity between conditions (x axis) and corresponding pvalue is depicted. **h)** Excerpt of motif similarity clustering as generated from TOBIAS based on binding site overlap. Motif 10 represents the de novo DENIS version of the Dux motif (see also (e)). **i-j)** TF-class assignment for two exemplary de novo motifs. The number of JASPAR motifs with high binding sequence overlap to the respective de novo motif is depicted as N, and the total number of motifs within the JASPAR database is given as DN. The plots show enrichment (x axis) for assigned TF-classes (y axis, one row per class). The enrichment is computed by counting the number of motifs in N assigned to a TF-class (depicted as n) and comparing it to the count of motifs in DN assigned to the same class (depicted as dn). Parts a) and c) are created with BioRender.com.

### Differential footprinting yields upregulated motifs

Next, we investigated the performance of the DENIS framework to predict genome-wide DNA binding events without any prior knowledge on the target motif. In practice, this is done by searching for UBMs without subtracting known motifs, and thus treating all FPs as part of the search space. This approach serves as an unbiased characterization of all DNA binding events from a single measurement, and can thus be compared between conditions. In the case of the Dux dataset, this approach can quantify differential binding events of novel motifs between Dux positive and Dux negative conditions, thus enabling us to identify condition specific upregulated motifs in an unbiased manner (Figure 3c).

The differential footprinting mode is realized by subtracting binding scores between conditions, in our example, the Dux negative from the Dux positive condition, which leaves us with the binding events unique to the Dux positive condition. Starting the framework on these positions and without a reference motif database, we received 178.034 Dux positive exclusive FPs. From these, DENIS reported a total of 90 UBMs, representing potentially 2C-like specific TFs. Of note, the motifs identified by DENIS should include both rediscovered database motifs as well as novel motifs. As expected, we found various levels of similarity to database motifs (Figure 3d-f, Supp. Figure 2c). These database motifs include NFYA^18^, SNAI2^19^, POU2F2^20^, KLF9^21^ and SP3^22^, which are well known to be active in developmental processes. Not surprisingly, we also identified the Dux motif as highly specific for the Dux positive condition. In addition to the 25 rediscovered motifs, we found 65 motifs that did not match the database, which are therefore considered novel.

Next, we asked whether the UBMs are indeed specific for the Dux positive condition. To this end, we ran TOBIAS^8^ with the newly discovered motifs joined with the JASPAR reference database. As expected, we found a significant enrichment for the newly discovered motifs to the Dux positive condition (Figure 3g), which supports their discovery from the Dux specific FPs. Interestingly, we saw an enrichment for both subgroups, namely the rediscovered and the novel motifs. We sought to further characterize the novel motifs by other means than motif similarity and used overlapping and clustering of binding sites, a method which enables grouping of novel and known motifs (Figure 3h). As previously shown^23–26^, TFs containing the same protein binding domain, but not necessarily the same motif, often act in a substituting or complementary manner. Consequently, enrichment of certain TF classes within clusters, derived from overlapping binding sites, can be used to assign UBMs to already annotated TF classes^27^. For our dataset, we utilized the top overlapping (>95% percentile) JASPAR motifs per UBM and checked their TF class annotations. Strikingly, for 50 out of the 90 discovered motifs we found a significant enrichment for a given TF-class (Supp. Figure 2d). Exemplary candidates include, among other, Basic helix-loop-helix factors (bHLH)^28^, C2H2 zinc finger factors^29^, T-Box factors^30^ and fork head/winged helix factors^31^, which are all well known to be important for developmental processes. In particular, our motif 10, which is assigned as the rediscovery of Dux, was enriched for Homeo domain factors, which Dux is a part of (Figure 3i).

Interestingly, one of the novel DENIS motifs showed a high enrichment for T-Box factors, a group of TF genes shared by all metazoan species (Figure 3j). This group of genes is well known to be involved in early embryonic development^30,32^.

Concluding, we have shown that our framework produces condition-specific motifs from footprinting data without prior knowledge. The UBMs were shown to have biological relevance by investigating their similarity to known motifs and assignment to TF-classes.

### Single-cell ATAC footprinting uncovers novel motifs in niche organisms

Most research in molecular biology is performed in a few standard model organisms for which a plethora of databases and background information are publicly available. However, aside from these, non-standard model organisms, defined as niche models^33^, lack these knowledge databases. Therefore, researchers working in these models are often forced to analyze their data based on comparison to standard organisms or have to translate their data by e.g. homology gene mapping.

In this context, DENIS is intended to work directly on niche model organism data, as it is able to identify UBM’s from footprints even if no motif database is provided. For illustration, we selected the zebrafish as a well known model organism with limited motif information in available databases. We used a scATAC-seq dataset^34^ dealing with hematopoiesis and the replenishment of blood cells. Existing cell type annotation for 12 cell types was used to aggregate single-cell signal into pseudobulk ATAC-seq data. DENIS was applied individually per cell type and without reference motif database (Figure 4a).

**Figure 4:**
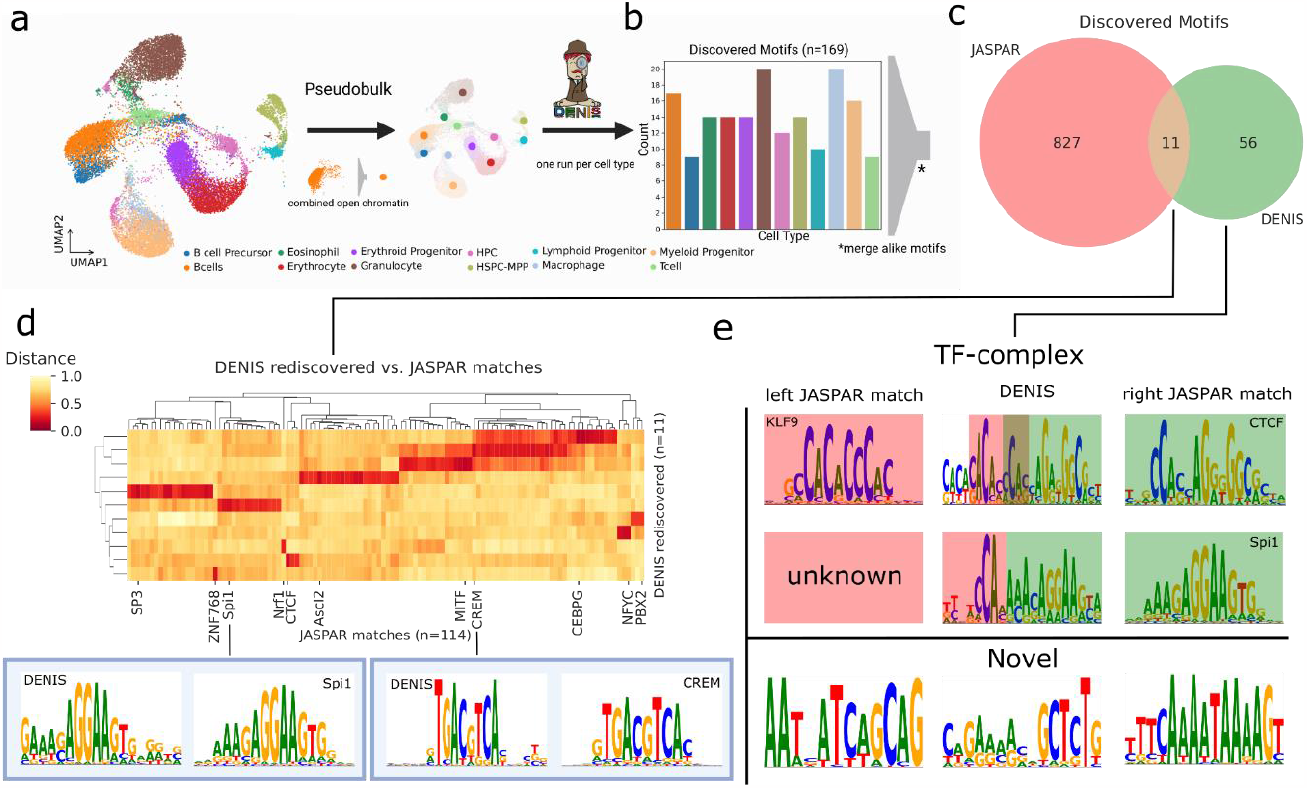
De novo motif generation within niche model organisms. **a)** Workflow to discover motifs per cell type. Initial scATAC-seq derived cell type clusters are merged into pseudobulks. DENIS de novo generates motifs on each cell type separately. **b)** Number of motifs per cell type. Motifs similar between cell types are considered active in multiple cell types and therefore merged. **c)** Overlap between JASPAR database and DENIS motifs. Out of 67 motifs discovered across all cell types eleven were found within the JASPAR database. **d)** Distance heatmap of DENIS rediscovered motifs (rows) to matches within the JASPAR database (columns). Column-labels show the closest JASPAR match per DENIS motif. Shown below are the DENIS motif (left) and JASPAR match (right) for Spi1 and CREM. **e)** Motifs with ambiguous or no match to the JASPAR database. Top are presumed TF-complexes. First row shows a combination of KLF9-CTCF, the second combines a novel motif with Spi1. The bottom part shows novel motifs created by the DENIS framework. Part a) and b) are created with BioRender.com.

First, we asked about the completeness of a de novo approach on this organism. In summary, we identified a total of 1.121.543 FPs across all cell types, with ∼70% containing at least one site used in motif generation (FPs may have more than one binding site). From these, DENIS was able to capture a total of 169 motifs (Figure 4b). As we expected some motifs, and thereby TFs, to be active in multiple cell types given, we performed a similarity distance comparison on all these motifs. This condensed the 169 initial motifs to 67 unique motifs (Supp. Figure 3a). Twenty-two of these motifs were found in two or more cell types (Supp. Figure 3b). This indicates that DENIS is able to find the same motifs independent of the distinct chromatin landscapes of different cell types. As ∼70% of human genes show orthologues to zebrafish genes, we first wanted to see how many of the DENIS motifs were found within motif databases^35^. Thus, JASPAR vertebrates were searched for motifs matching the discovered UBMs. Eleven motifs could be assigned to database motifs. When checking the corresponding human TF sequences, Ensembl^36^ reported eight of the eleven factors to have a highly conserved homologue within zebrafish (Supp. Figure 3c), with the GeneCards compendium^37^ assigning homologs to the remaining factors (PBX2, CEBPG and ZNF768). Furthermore, many were found to play a role during hematopoiesis, e.g. SP3^38^, Spi1^39^, CTCF^40^ and PBX2^41^ (Figure 4c-d, Supp. Figure 3d), supporting our hypothesis on rediscovering active TF by rediscovering TF motifs.

Next, we asked about the other 56 motifs that were not assigned to a given JASPAR motif. Upon visual inspection, we classified them into two groups, namely i) extended motifs, and ii) pure novel motifs. While the first group is the smaller subgroup with about ⅓ of the motifs, it is characterized by partial similarity to one or multiple database motifs. As shown in (Figure 4e top, Supp. Figure 3e), these motifs are potential TF complex motifs, with both parts to be known single motifs. An exemplary case is shown for a DENIS motif that matches KLF9 and CTCF, both assigned to have a protective role against DNA methylation^42^. Other examples were found with one of the parts to be a known database motif, while the second part is novel. The second subgroup of motifs is not related to JASPAR motifs (Figure 4e bottom, Supp. Figure 3f). We consider this presumed overlap of novel and established motifs to further support the validity of motifs generated by our framework.

Finally, we checked the distribution of novel and rediscovered motif binding sites in FPs utilized to create UBMs. We found that nearly all of the utilized FPs (∼94%) contained a binding site associated with a novel motif, whereas only ∼43% FPs contained binding sites of rediscovered motifs, this stresses the need for an organism agnostic motif generation pipeline such as DENIS.

### DENIS uncovers cell type specific motifs

So far, we identified motifs in the WT condition of each cell type^34^. However, the dataset also provides a complementary *gata2b* knockout (KO) condition, enabling us to *de novo* analyze key differences between two hematopoiesis conditions. Thus, we executed DENIS in differential footprinting mode to yield motifs specific to the WT condition. The framework discovered 57 motifs across all cell types, and as before, consensus motifs were created to accommodate motifs occurring in multiple cell types, resulting in ten motifs with FPs unique to the WT condition (Figure 5a). Comparing these motifs to JASPAR, DENIS classified six as rediscovered and the remaining four as novel (Figure 5b-c).

**Figure 5:**
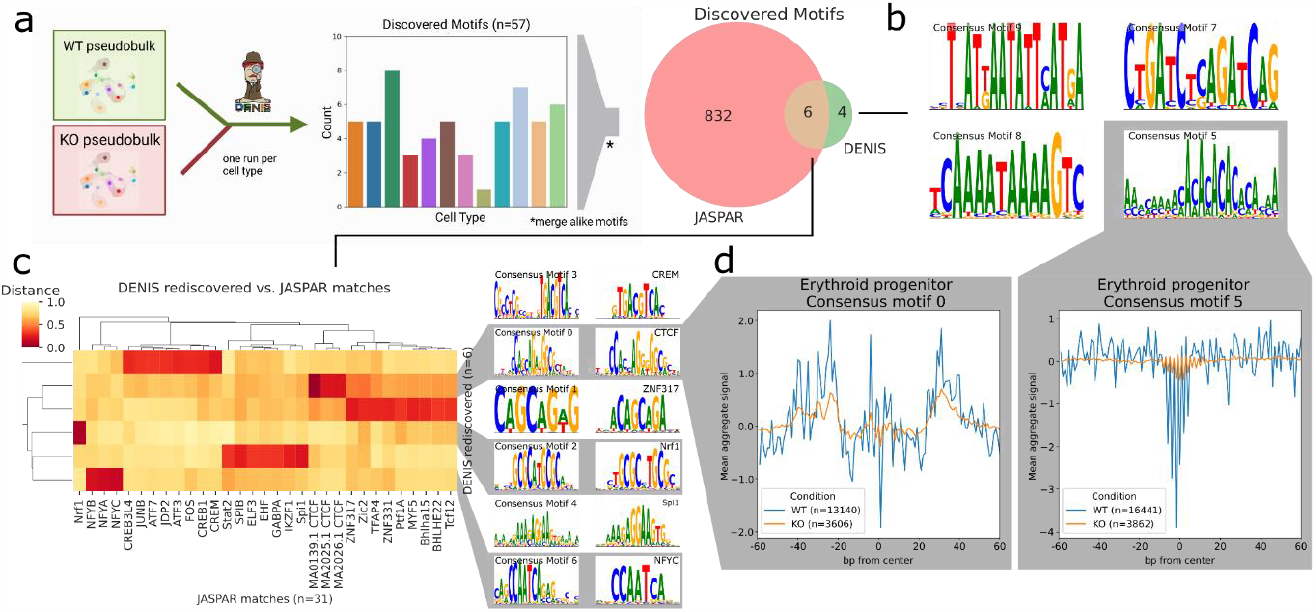
Differential motif generation and footprinting. **a)** Workflow to unravel wildtype (WT) exclusive motifs. Pseudobulks of WT and knockout (KO) per cell type are provided to DENIS differential footprinting mode. Similar motifs across cell types are merged. Rediscovered motifs are identified with the JASPAR database. Created with BioRender.com. **b)** Novel DENIS motifs identified to be unique for the WT condition. **c)** Distance heatmap of DENIS to JASPAR motifs with de novo motifs in rows and similar factors as columns. The two motif logo columns on the right show de novo DENIS motifs and their closest JASPAR match. **d)** Two aggregated footprints for WT vs. KO (orange and blue line) in erythroid progenitor cells including number of binding sites (n). Plots indicate a motif centered view with a width of 120bp.

We compared the rediscovered motifs and their originating cell types to WT enriched motifs described by Avagyan et al.^34^. One interesting motif here is consensus motif 0, which is almost an exact replica of CTCF and found by our framework in all cell types except for granulocytes. Interestingly, Avagyan et al.^34^ classified CTCF in more detail and reported it to be highly specific in the WT condition of five selected cell types, to be moderately specific in another seven cell types, and granulocytes to completely lack CTCF activity. Our results completely support the findings of Avagyan et al.^34^, and more importantly, highlight the capability of DENIS to robustly detect motifs, even if the TF signal considerably varies between the cell types investigated.

Another example is consensus motif 4. It was found in B cell precursor, B cell, Granulocyte, Macrophage, Myeloid progenitor and Eosinophils. This UBM can be assigned to a group of seven JASPAR motifs (Figure 5c), of which SPI1, SPIB, Stat2, EHF and GABPA were found to be enriched in the dataset by Avagyan et al.^34^ as well.

Furthermore, we asked whether all UBMs (rediscovered and novel) exhibit steric hindering of Tn5 transposase activity, as shown by aggregated footprinting plots. By plotting Tn5 signals centered at motifs, we found six out of ten DENIS motifs with a strongly aggregated footprint signal (Supp. Figure 4). We saw a clear footprint for consensus motif 0, which is not surprising, as the matching TF CTCF is well-known to create a strong footprint signal due to its role as a chromatin remodeler. However, we also identified this footprint to be stronger in WT in comparison to KO, which supports the assumption that this TF is enriched in WT footprints (Figure 5d left). In addition, we found strong WT specific footprints for consensus motif 5, a novel motif, further proving evidence of TF activity to be co-localized with the DENIS motifs (Figure 5d right). Additional FPs were found for consensus motif 2 (Nrf1), consensus motif 3 (CREB3L4, JUNB, ATF7, JDP2, ATF3, FOS, CREB1, CREM), and consensus motif 4, also suggesting these TFs to be highly active (Supp. Figure 4). Interestingly, consensus motif 6, partially rediscovered as the NF-Y TF, shows no FP but periodic DNA blocking (Supp. Figure 4f). NF-Y is reported to be essential for the replenishment of hematopoietic stem cells^18^. We hypothesize the periodic pattern to reflect its three subunits binding together (NFYA, NFYB, NFYC).

Concluding, identified motifs are shown to be specific for condition and cell type. Furthermore, most motifs are supported by cell type specific aggregated footprints for rediscovered and novel motifs. These findings render DENIS especially applicable for motif analysis across many samples or cell types resulting from bulk ATAC-seq or scATAC-seq.

## Discussion

The utilization of DNA binding motifs as a surrogate for TF binding and their role in predicting gene regulation has been investigated for decades. However, even for human, arguably the most characterized organism, less than 50% of TFs have been described by a TF binding motif (JASPAR CORE), hindering unbiased genome-wide studies on TF binding and TF activity. Consequently, these motif based approaches fall short when analyzing niche model organisms, as the rate of available motifs in such cases is vanishingly low.

DENIS utilizes a completely new approach for de novo motif generation from ATAC-seq data. Briefly, DENIS can find UBMs from complex mixtures of sequences by using an iterative logic to aggregate motifs from FPs that are represented with different frequencies in the data. While production of UBMs can be a challenge, downstream analysis and assessment of functional significance remain just as critical. In this manuscript we introduced a series of methods, such as structured comparison to motif databases, or the overlap of binding site locations that enable the characterization of UBMs. DENIS defines a new class of computational tools and strategies to filter candidates towards meaningful results.

The overall ability to find UBMs is highly dependent on the cell type and organism investigated. For example, when analyzing the Dux TF data, we noticed a strong difference in the number of generated motifs between the Dux positive and Dux negative condition (48 vs. 3 motifs). We attribute this disparity to the relatively unexplored mechanisms of epigenetic regulation during zygotic genome activation. Many factors that are active at this stage have never been characterized by ChIP-seq analysis. This assumption is supported by the massive change of the overall chromatin structure in this dataset^8^. In contrast, the Dux negative condition constitutes a well studied WT mESC that is not expected to hold many novel active TF candidates. This was further confirmed during the differential analysis of the Dux TF data. A characterization of Dux positive *de novo* motifs by binding site overlap showed our rediscovered Dux motif to cluster closely with Dux and other Homeo domain factors. In line with control factor Dux, we found most UBMs to show a TF-class enrichment. An illustrative example is motif 37 enriched for the T-Box factor class that lists 17 known associated genes^30^, of which 15 are found in the JASPAR database. The two remaining genes TBX10 and TBX22 are not present in JASPAR, allowing speculation whether the novel motif predicted by DENIS is the binding motif of either one of these genes or a TF that has yet to be described, but showing high overlap with the T-Box factor class.

When applying DENIS to a niche model organism situation with zebrafish scATAC-seq, a comparable low motif rediscovery rate is found. Of 67 UBMs identified from the WT cell samples, only 11 could be assigned to a known human motif in JASPAR’s vertebrates collection, which does not contain any motifs derived from zebrafish. Interestingly, the respective genes to all 11 candidates were found to be highly conserved between zebrafish and human. Therefore, we assume binding motifs of these factors to be conserved as well. However, for the assignment of the remaining motifs, pure motif similarity falls short, potentially because the TFs have diverged too much and as such are too dissimilar to assign. This is also evident in the example of WT vs. *gata2b* KO, where we were not able to identify a motif with significant similarity to a given GATA2 within JASPAR, even though this would be expected based on our differential approach to collect FPs exclusive for the WT condition. We hypothesize that zebrafish’s *gata2b* is part of the novel UBMs but possesses a binding motif too different from given GATA2 and thus can not be assigned. Regardless of the UBMs affiliation to either be a novel or rediscovered motif, we found a considerable amount of motifs in more than one cell type or even subsets of cell types. These are assumed to originate from the same TF active in different environments and roles, which our approach is able to identify despite drastically changing chromatin landscapes (available FPs) between cell types. These findings support our claim for biological significance of the UBMs.

In summary, DENIS is the first versatile framework for unbiased TF activity analysis via *de novo* motif generation and binding site assignment based on ATAC-seq footprinting. DENIS constitutes a versatile and cost effective tool when screening for a binding pattern of a factor that refuses detection by e.g. ChIP-seq. However, as ATAC-seq footprinting is only able to report a binding event, but not the specific protein bound, we want to emphasize that extended methods such as mass spectrometry are needed to further investigate promising motif candidates and corresponding protein assignments. Nevertheless, the ability of DENIS to run without a reference motif database generates opportunities for model and niche model organisms and enables detection of highly promising candidates for wet-lab validation or to significantly improve coverage of motif databases.

## Data availability

Dux overexpression ATAC-seq, and ChIP-seq data are available from GEO under the accession GSE85632. Hematopoiesis scATAC-seq data is available from GEO under the accession GSE151232.

## Methods

### Data

#### Benchmark of motif generation tools

Motif binding sites were collected from JASPAR^6^. Datasets with a binding site count from 100 to 15000 were created for one motif (MA0474.1) and four motifs (MA0474.1, MA0035.3, MA0461.1 and MA0480.1). Binding sites of the latter were divided into a fixed ratio (40, 30, 20 and 10%). The datasets were finally used to run selected motif discovery tools (MEME, DREME, GLAM2) and evaluate their output.

#### Benchmark motif selection

A dataset of 20 ChIP-seq motifs was constructed to test MEMEs ability to find motifs in a heterogeneous environment similar to ATAC-seq data. Binding sites of motifs were collected from JASPAR and flanking regions were added until each sequence had a total length of 100bp. The number of motif binding sites ranged from 395 to 322.803 with the dataset being comprised of 659.078 binding sites total.

#### Exemplary bulk ATAC-seq data

Public ATAC-seq data of mouse embryonic stem cells was obtained from (Hendrickson et al., 2017) The data comprises Dux-positive and -negative samples, which code for Dux overexpression and Dux wildtype, respectively. The data was downloaded and then prepared for analysis using TOBIAS^8^. ATAC-signal was corrected for the Tn5 transposase bias and converted to an FP binding score. The FP score was used to create and analyze binding motifs using DENIS.

To test DENIS ability to rediscover Dux, motifs from JASPAR2022 core vertebrates and HOCOMOCOv11 core human and mouse were combined. Next, all motifs similar to Dux are removed from the dataset to ensure Dux binding sites to be treated as novel binding sites. This was done by computing a distance score using TOBIAS. All motifs below a distance threshold (<0.4) were removed, which resulted in the removal of 38 out of 1595 motifs. The remaining motifs were then provided as a database during Dux rediscovery.

#### Exemplary single-cell ATAC-seq data

Public scATAC data of zebrafish hematopoietic cells was taken from (Avagyan et al., 2021). The data comprises hematopoietic cell types over a wildtype vs. knockout condition. The data was downloaded and preprocessed by aligning reads using STAR^44^, followed by quality filtering using EpiScanpy^45^. Remaining cells were assigned to cell types using cell type annotation as provided by^34^. Thereafter, pseudobulks were created by combining reads of cells assigned to the same cell type. This enabled data preparation with TOBIAS^8^ (see above) followed by DENIS analysis on each cell type.

### DENIS Framework and Code availability

The DENIS framework is publicly available from https://github.com/loosolab/denis and is split into three parts (see below). The first part performs i) footprinting analysis and footprint characterization (known/unknown), in which DENIS identifies protein binding locations in a genome-wide manner. Second, unknown FP sequences are extracted and are used to create novel motifs. Finally, in iii) novel motifs are provided to downstream analysis modules. The exact steps of a DENIS framework are dependent on the supplied data. An exemplary framework run is given via the repository.

#### i) Footprinting analysis

This step will identify FPs by scanning a continuous binding score, as produced by a footprinting tool such as TOBIAS in Bigwig format^8^, by calling peaks. A peak is considered a FP if it meets certain parameters. It has to have a flat top, be within a defined width and has to exceed a local height limit. FPs in close proximity will be merged if the gap between them is below a given width and depth threshold. Furthermore, assuming a motif database is available, FPs are scanned for overlap to motifs. Regions where overlap is detected are removed, filtering the FPs to regions without a known binding TF.

If two scorings (e.g. two conditions) are given, the scores are combined to a differential FP score prior to identifying the FPs. The differential score is computed by subtracting the second from the first score. Importantly, values below zero are set to zero. This results in a differential binding score, representing regions unique to the first score, hence FPs unique to the first condition.

#### ii) De novo motif discovery

Novel motifs are generated using FPs identified in the prior step. This is accomplished using the MEME tool in consecutive runs with decreasing amount of FPs. After each MEME run, significant motifs are filtered based on e-value. The filter threshold is dynamically chosen by fitting the e-values to a function, on which a knee location algorithm is applied. Afterwards, FPs used to create significant motifs are filtered, by removing the exact motif location within each FP, retaining the flanking parts of the FP for the following iteration. Keeping the flanking parts is done to identify motifs of closely bound TFs. The reduced FP set is then used to further create novel motifs. This cycle repeats until no more motifs are generated or no significant motifs are found for several iterations. Finally, all de novo generated motifs are clustered and similar motifs are merged. The clustering is done utilizing TOBIAS with a user defined distance threshold (throughout this work set to 0.4 for all merging operations), below which motifs are combined. The framework proceeds with the resulting consensus motifs.

#### iii) Downstream analysis modules

Finally, novel motifs are subjected to several types of analysis. Given a motif database, a set of canonical motifs, which is either automatically chosen by the framework or defined by the user, is analyzed alongside the novel motifs. Each motif within this set is used to scan for binding sites throughout the genome. The binding sites are used to investigate their enrichment in open chromatin as well as to annotate the ones within open chromatin to genomic features in proximity. The latter requires genomic feature information being supplied to the framework. Binding site annotation of each motif is then followed by feature enrichment analysis. Assuming provided feature information contains genes, a genes set enrichment analysis is computed, showing biological processes, cellular components and molecular functions related to each motif. Additionally, motifs are compared to provided database motifs and the most similar database motifs are shown for each.

## Acknowledgements

This study was funded by the German Research Foundation (DFG):EXC2026/1, KFO309 and LOEWE iCANx to M.L, as well as the Max Planck Society. We would like to thank Carsten Kuenne for critically reviewing the manuscript.

## Author contributions

M.L conceived the study. H.S. and M.B. designed the software, H.S. implemented the software. H.S., M.B., V.H. and M.L. preprocessed and analyzed data. H.S., M.B. V.H. and M.L. wrote the manuscript. M.L. supervised the project.

