## Supplementary Figures for "Uncovering uncharacterized binding of transcription factors from ATAC-seq footprinting data"

---

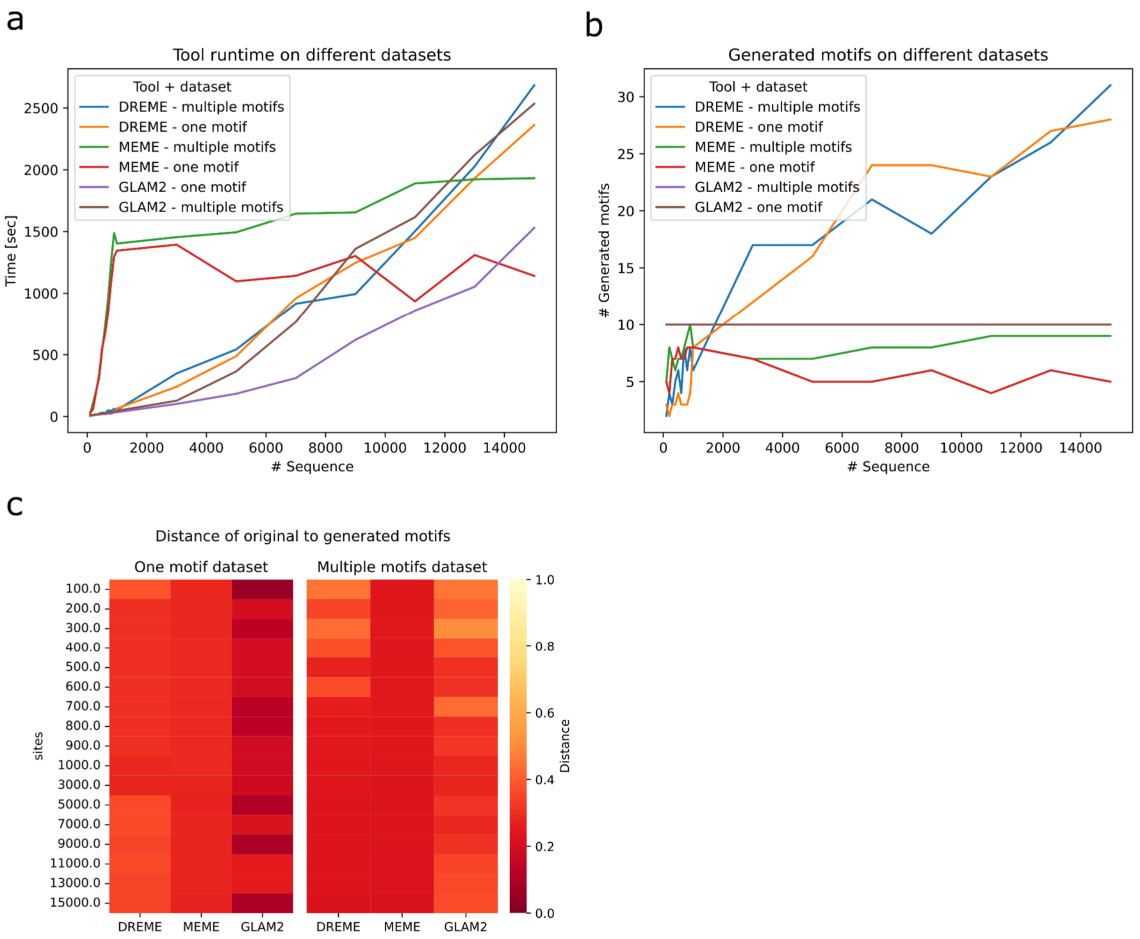

**Supplementary Figure 1: Comparison of motif discovery tools**

*a) Tool runtime in dependence of number of supplied sequences. Each tool was tested on a dataset containing one motif and a dataset with multiple (four motifs).*
*b) Number of generated motifs in dependence of supplied sequences on the same datasets as in a). GLAM2 is not able to infer the number of motifs based on the data and therefore always generates ten motifs.*
*c) Distance of the closest output motif to the original motif (mean distance for multi-motif datasets). Rows show the number of sites of the given dataset columns show the motif tool. The left side shows runs on datasets with one motif the right are runs on datasets with multiple motifs.*

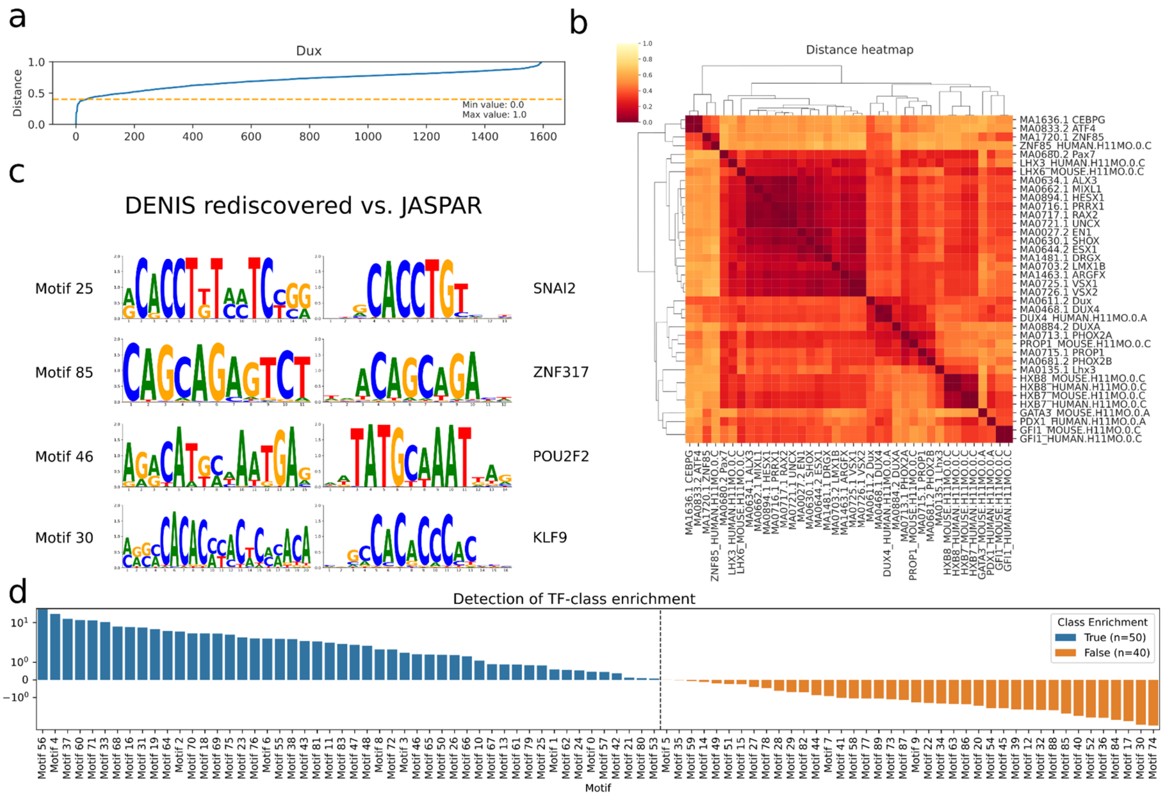

**Supplementary Figure 2: Dux dataset analysis**

*a) All database motifs sorted by their distance to Dux. Motifs below the threshold are removed for the Dux rediscovery analysis.*
*b) Heatmap of the distances between all motifs below the threshold in a).*
*c) DENIS motifs (left) generated during the Dux differential analysis compared to the closest match of the JASPAR database.*
*d) Categorization of DENIS motifs to show enrichment for TF-classes. Based on the difference between maximum odds ratio and Q3 value (interquartile range) per motif.*

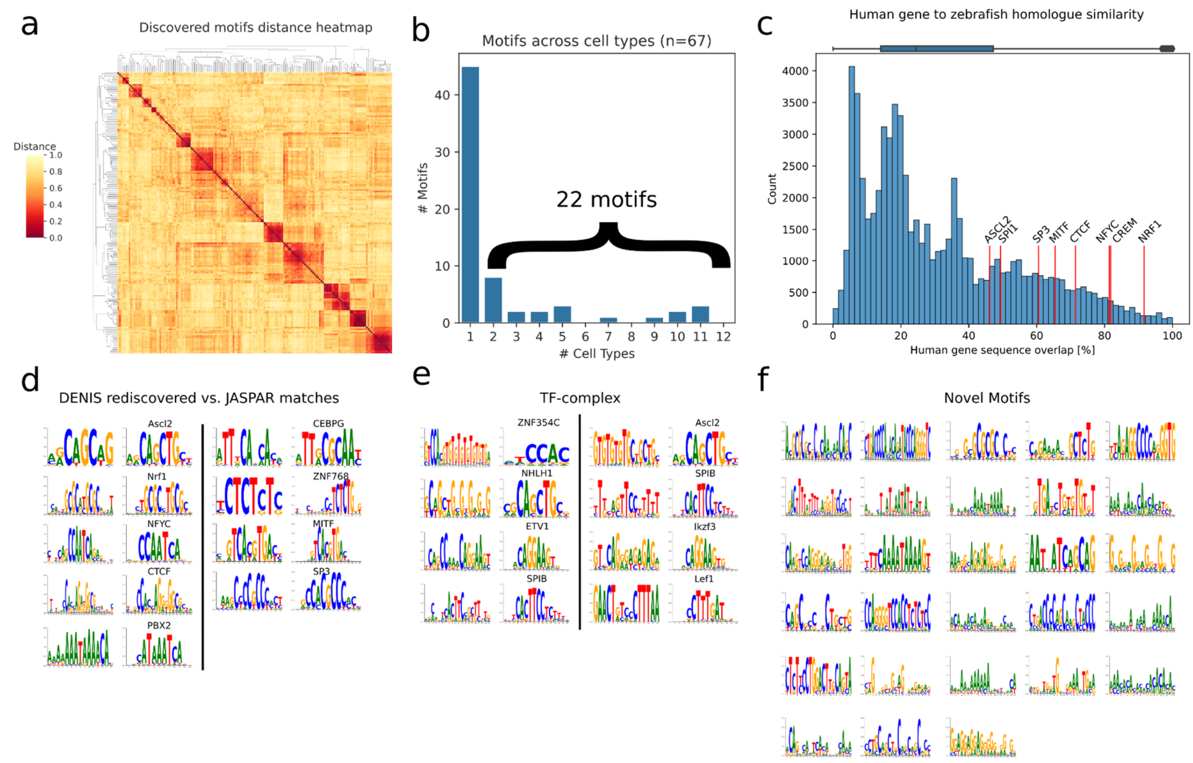

**Supplementary Figure 3: Zebrafish dataset analysis.**

*a) Distance of DENIS motifs discovered during the WT cell type runs to each other. Motif clusters are combined as they are discovered in multiple cell types.*
*b) The number of motifs counted for their appearance in cell types.*
*c) Distribution of percent sequence overlap of human genes to their zebrafish homologs. The motif of marked genes is rediscovered by the DENIS framework.*
*d-f) Motifs created by the DENIS framework during the zebrafish WT analysis. d) are rediscovered DENIS motifs (left) and their closest JASPAR match (right). e) are assumed to be TF-complex motifs (left) where only part of the motif is known accompanied by their best JASPAR match (right). f) are novel DENIS motifs that did not match a JASPAR motif.*

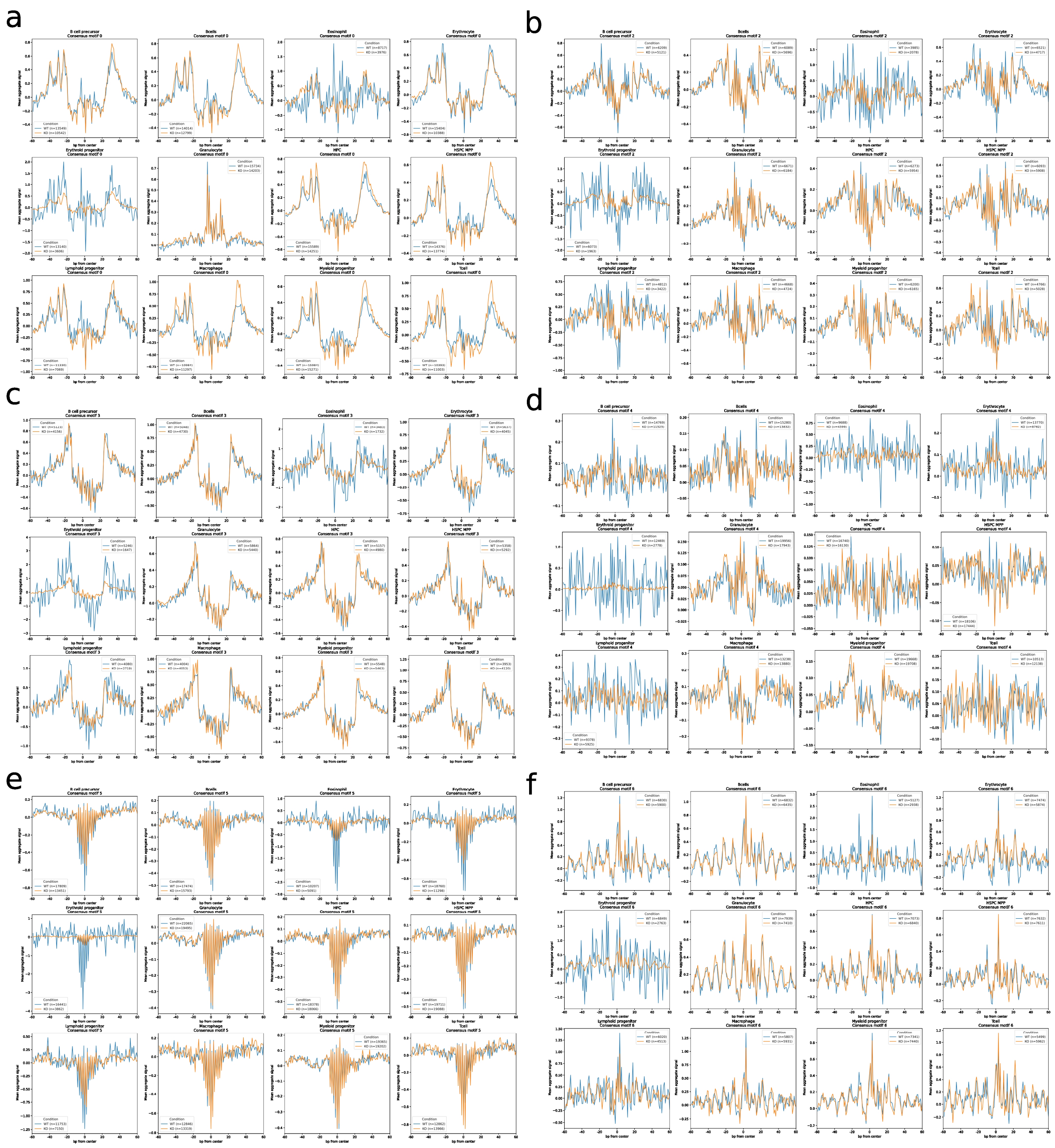

**Supplementary Figure 4: Aggregated FPs.**

*Aggregated FP of WT (blue) vs. KO (orange) motif positions (created with TOBIAS PlotAggregate) each plot shows the FP for one of the cell types. FPs correspond to consensus motif 0 a), consensus motif 2 b), consensus motif 3 c), consensus motif 4 d), consensus motif 5 e) and consensus motif 6 f).*
